# Cell-state conditioning in AlphaGenome 2 reproduces an unseen interferon response but shows no reliable stimulus-specific eQTL effects

**DOI:** 10.64898/2026.11.16.767041

**Authors:** Haodong He, Jianxuan Sun, Jingyu Xu, Maimaiti Abudureyimu, Yuxuan Yang, Yifan Huang, Zhihua Wang, Qidong Xia, Shaogang Wang

## Abstract

**Background:** Sequence-to-function models such as AlphaGenome 2 can be conditioned on a cell state supplied as a single-cell expression profile. Whether this conditioning captures the effects of an experimental stimulus, on gene expression or on the activity of regulatory variants, has not been tested.

**Results:** We conditioned AlphaGenome 2 on interferon-β-stimulated and unstimulated CD14+ monocytes from a published single-cell dataset, using cells that were not used to measure the response. Predicted and measured expression changes agreed across 119 genes (Spearman ρ = 0.615; 95% CI 0.486 to 0.716), and agreement was essentially unchanged when these genes were removed from the conditioning input (ρ = 0.607). Across fifteen conditioning states, the predicted response scaled with each state’s measured similarity to the interferon response (slope 0.73, Pearson 0.92). We then asked whether the same conditioning identifies stimulus-specific expression quantitative trait loci (eQTL) in two monocyte stimulation studies. The main comparison in our analysis plan, on interferon-γ-stimulated monocytes, was inconclusive (p = 0.096). Of four replication arms, only influenza infection separated activation-specific from resting-specific eQTL (p = 0.0019; 0.0076 corrected for four arms); the difference arose from near-zero predicted effects in the resting condition and disappeared when both conditions were required to have measurable predicted effects (p = 0.443). Two analysis issues are likely to affect similar evaluations: 71.5% and 75.9% of eQTL labelled condition-specific by credible-set membership had a significant effect in the other condition, and a ratio statistic filtered on the larger of its two terms converted a baseline difference into apparent condition dependence.

**Conclusions:** Conditioning on a single-cell profile transferred the expression response to an unseen stimulus, but we found no reliable evidence that it transfers stimulus-dependent variant effects. Evaluations of conditioned models should test these two capabilities separately, define condition-specific eQTL by testing both conditions, and require measurable effects in both conditions before comparing them.

## Background

Sequence-to-function models predict regulatory activity, such as chromatin accessibility and gene expression, directly from DNA sequence [1, 2]. Until recently, these models predicted a fixed set of cell types and assays, namely those represented in their training data. Two models released in late 2026 instead condition their predictions on a cell state supplied by the user as single-cell expression data, so that predictions can be requested for states outside the training catalogue [3, 4].

Many of the questions that motivate such predictions concern perturbed states: the response of a cell to a cytokine, a pathogen or a drug, and the regulatory variants whose effects appear only after stimulation. Studies of response eQTL in immune cells have shown that a substantial fraction of genetic effects on gene expression are modified by stimulation [5–8]. Reference catalogues, by contrast, are built largely from unperturbed tissue, so a user interested in a perturbed state has to supply it.

Predicting transcriptional responses to perturbations from single-cell data is an established problem. Generative models such as scGen, CPA and GEARS predict responses for unseen cell types or perturbations [9–11], and scGen used the interferon-β-stimulated blood cells analysed here as a benchmark. Recent benchmarks show that simple linear baselines perform as well as deep learning models at predicting the effects of genetic perturbations [12]. Conditioned sequence models differ from these approaches in that they predict expression from DNA sequence given a state, and can therefore also be used to predict how sequence variants act in that state. Sequence models have so far explained differences in expression between individuals poorly [13, 14]. Independent evaluations of the first-generation AlphaGenome found that it registers in-silico promoter disruption locally but often fails to propagate it to downstream exons [15], and that annotations derived from it do not improve GWAS locus discovery beyond standard functional annotations [16]. None has evaluated cell-state conditioning.

Two capabilities need to be distinguished. A conditioned model may reproduce how expression differs between two states without reproducing how the effects of individual variants differ between them, because the condition is a representation learned from expression profiles, which describe how a state looks on average rather than how it changes the consequences of individual variants.

Here we test both capabilities in AlphaGenome 2 using interferon-β-stimulated CD14+ monocytes, a stimulated state that is not among the model’s built-in conditions. We ask whether the predicted expression response tracks the measured response, how the predicted response scales across fifteen conditioning states, and whether the same conditioning identifies stimulus-specific eQTL in two monocyte stimulation studies. We also describe what the built-in conditions of both model generations contain, and two analysis issues that produced a spurious result in our site-level analysis and are likely to affect similar evaluations.

## Results

### Predicted expression response to interferon-β

We conditioned AlphaGenome 2 on interferon-β-stimulated and unstimulated CD14+ monocytes from a published single-cell experiment [17] and compared the predicted log2 fold change between the two states with the measured one. Each condition was built from 256 cells, the maximum accepted by the encoder, drawn once per library without replacement. The measured response was computed from the remaining 2,529 unstimulated and 2,334 stimulated CD14+ singlets, pooled to libraries of 5.63 and 5.76 million counts, so that no cell contributed to both the condition and the measurement. Of the 246 genes in the 40 input intervals that had a measured value, 119 passed the activity filter (maximum predicted activity above 1) and form the analysis set (Methods).

Predicted and measured log2 fold changes agreed at Spearman ρ = 0.615 across the 119 genes (window-clustered bootstrap 95% CI 0.486 to 0.716; 35 windows of 1 Mb; Fig. 1a). Removing the ten genes with the largest absolute measured log2 fold change left ρ = 0.563 (n = 109; 95% CI 0.419 to 0.674), so the agreement was not driven by a few strongly responding genes. Deleting all 119 analysis genes from the conditioning input, which reduced it from 35,635 to 35,516 genes, gave ρ = 0.607 with the measured response, and these predictions correlated with the original ones at ρ = 0.989 (Fig. 1b, c). The predictions for these genes therefore did not depend on their own expression values in the conditioning input.

**Figure 1:**
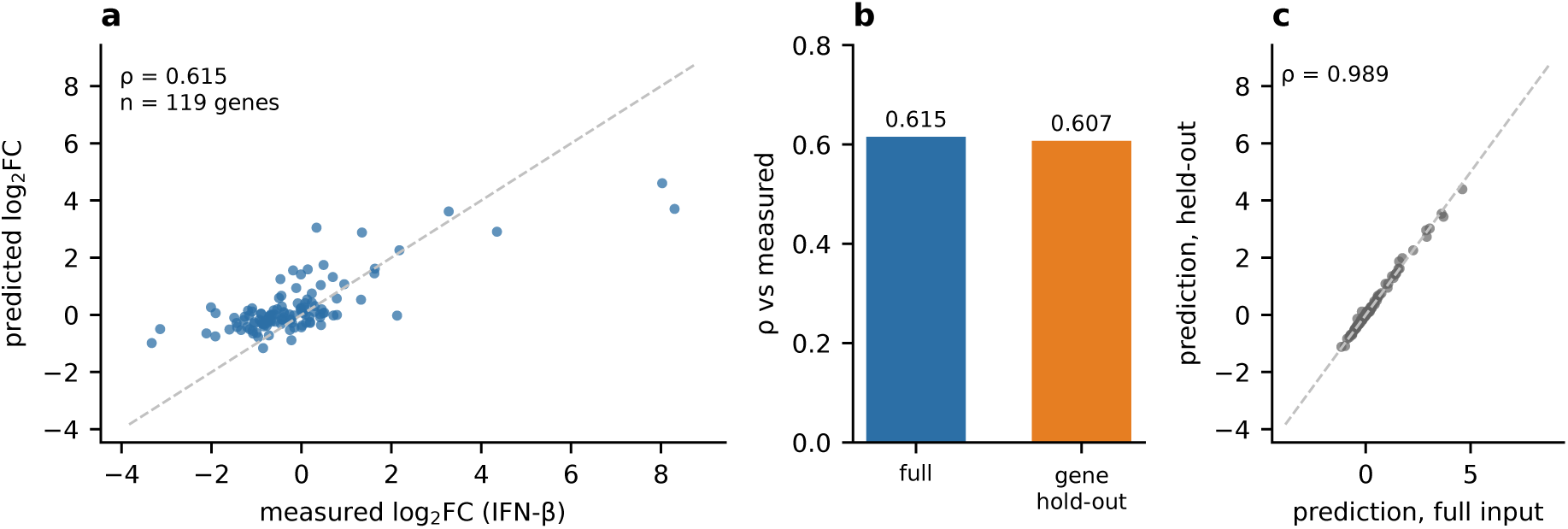
Expression-level prediction and gene hold-out. (a) Predicted versus measured log2 fold change for the 119 analysis genes under the matched interferon-β condition (Spearman ρ = 0.615; window-clustered bootstrap 95% CI 0.486–0.716). (b) The same comparison after all 119 analysis genes were removed from the conditioning input (ρ = 0.607). (c) Agreement between the original and hold-out predictions (ρ = 0.989).

Table 1 summarises the supporting analyses. In our initial analysis, the measured response was computed from the same 256 cells that built each condition, which gave ρ = 0.486 at a library depth of 557,000 counts. Binomially thinning the disjoint counts to that depth gave ρ = 0.434 ± 0.053 over 20 draws. Sharing cells therefore raised the estimate by about 0.05, within the standard deviation of the thinning, whereas the lower depth accounted for about 0.18 of the difference (Additional file 1, Fig. S1). In a split-half null, the control cells were divided into two halves that served as the two conditions; the mean correlation was +0.027 (SD 0.229), and none of 50 draws reached the observed value, which gives the smallest attainable one-sided p of 1/51 = 0.020 (Additional file 1, Fig. S2a). Because the split-half predictions are about an order of magnitude smaller than the real ones, this null shows that the pipeline does not create correlation from noise, but not whether an unrelated condition could reach the same value; the calibration below addresses that question.

**Table 1.** Supporting analyses for the expression-level comparison.

| Analysis | Statistic | Value |
|---|---|---|
| Main comparison | Spearman ρ, 119 genes | 0.615 (95% CI 0.486–0.716) |
| Target genes held out of the conditioning input | Spearman ρ | 0.607 |
| Held-out predictions vs original predictions | Spearman ρ | 0.989 |
| Depth-matched disjoint estimate | Spearman ρ, 20 thinnings | 0.434 ± 0.053 |
| Split-half null (control cells halved, 50 draws) | mean ρ, SD | +0.027, 0.229 |
| Split-half null, draws at or above observed | count | 0 of 50 (one-sided p = 0.0196) |

The conditioning and measurement cells were disjoint but came from the same eight donors. Because demuxlet assigns every cell to one of the eight donors, a donor-disjoint split is possible; it was not run.

### Calibration across fifteen conditioning states

To ask whether the predicted response depends on how similar the conditioning state is to the target state, we conditioned the model on fifteen states: seven unstimulated cell populations, the same seven after interferon-β stimulation, and the matched interferon-β monocyte state. For each state, the measured overlap is the Spearman correlation between that state’s measured log ratio relative to unstimulated CD14+ monocytes and the measured interferon-β response; the model value is the same correlation computed with the predicted log ratio after conditioning on that state (Methods). The matched condition has a measured overlap of 1 by construction.

The model value was linearly related to the measured overlap, y = 0.728x − 0.025 (Pearson 0.923, Spearman 0.850; Fig. 2). Correcting for measurement error in x, whose reliability was 0.983, gave a slope of 0.741, and the intercept was indistinguishable from zero. On average, the model therefore recovered about three quarters of the rank structure that a state’s measured contrast shared with the target response. Recomputing the measured axis against the CD14+ monocyte baseline that excludes the conditioning cells changed the slope from 0.728 to 0.731 and the corrected slope from 0.741 to 0.743.

**Figure 2:**
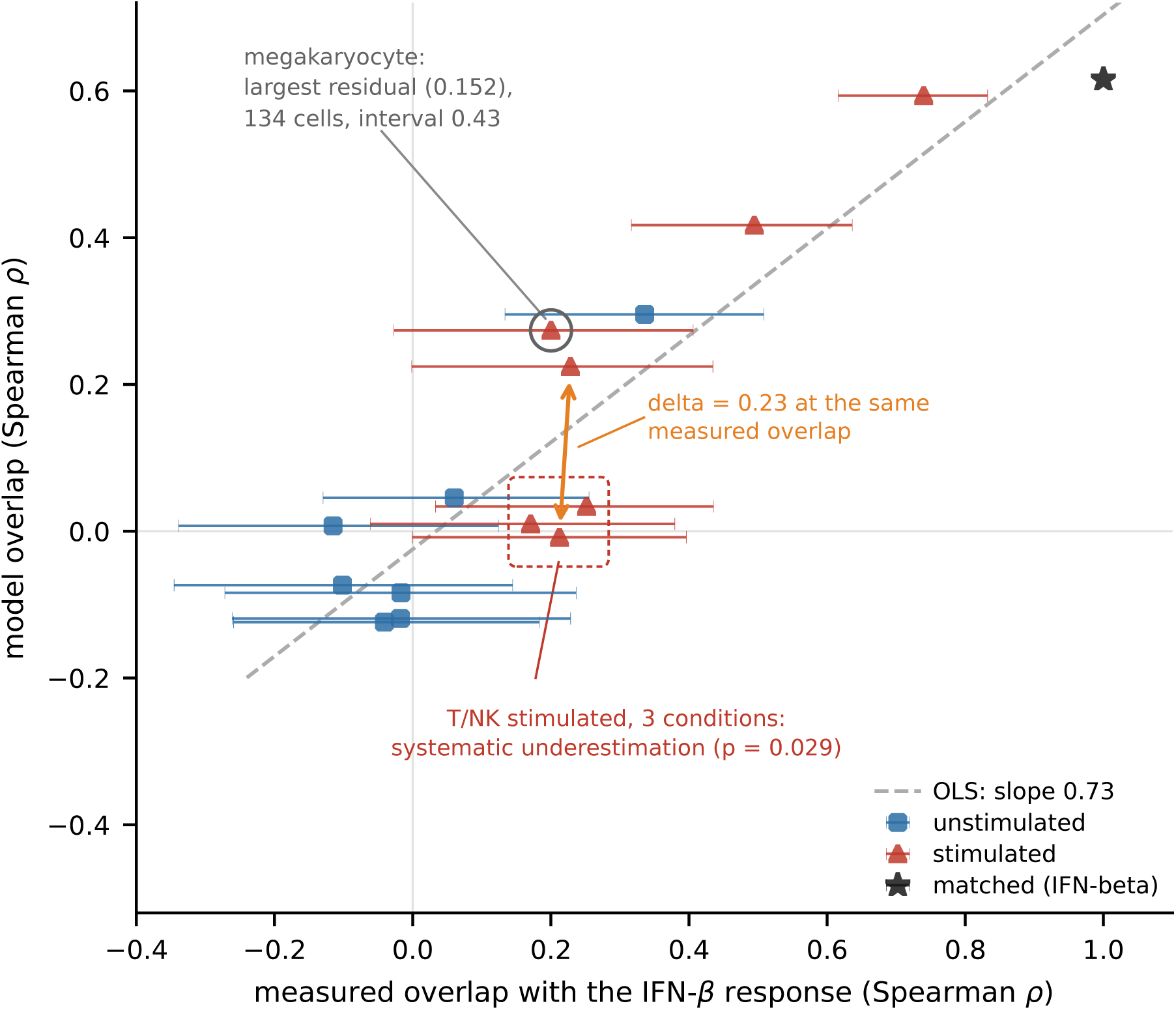
Calibration of predicted against measured cross-condition overlap. Each point is one of 15 conditioning states: seven unstimulated cell types (circles), the same seven after IFN-β stimulation (triangles), and the matched IFN-β monocyte condition (star). The x axis is the measured overlap between that state and the monocyte IFN-β response, that is, the Spearman correlation across the 119 analysis genes between log2 expression of the state relative to unstimulated CD14+ monocytes and the measured IFN-β log2 fold change; horizontal bars are 95% window-clustered bootstrap intervals. The y axis is the same quantity computed from model predictions. Dashed line: ordinary least squares fit (slope 0.728, intercept −0.025; Pearson 0.923, Spearman 0.850; 14-point fit without the matched condition, slope 0.826). Three features are annotated: the three stimulated T/NK states carry the three negative residuals (permutation p = 0.029 for that partition); the CD8 T and B cell conditions have almost identical measured overlap (0.212 and 0.228) yet differ by 0.23 in the model, so the residual is not a smooth function of the x axis; and the stimulated megakaryocyte condition has the largest residual (0.152) and rests on the fewest cells (134 cells; interval width 0.43). *Measured baselines.* The x axis uses all singlet cells of each annotated cell type (2,785 CD14+ monocytes in the control library, 5,161 CD4 T, 1,082 CD8 T, 916 NK, 1,307 B, 203 dendritic, 122 megakaryocyte, 739 FCGR3A+ monocytes), with no cells excluded; the measured response on the y axis uses CD14+ monocytes excluding the 256 cells per library that built the conditioning input (2,529 control, 2,334 stimulated). The two axes therefore use different cell sets (Methods; Additional file 1, Table S2).

Unstimulated FCGR3A+ (CD16+) monocytes, the population closest to the target, gave a model value of 0.295, about half the matched condition’s 0.615, at a measured overlap of 0.336 (95% CI 0.133 to 0.508). The fitted line predicts 0.22 at this overlap, so the point lies 0.075 above the line: the contrast between the two monocyte populations shares structure with the interferon response, and the model reproduced most of it. The six other unstimulated conditions received model values between −0.12 and +0.05.

Deviations from the line were associated with lineage. Stimulated CD8 T and B cells had similar measured overlaps (0.212 and 0.228) but different model values (−0.008 and +0.224), a gap of about 0.23. Among the seven stimulated conditions, the three T and NK populations had the only negative residuals (CD4 T −0.090, CD8 T −0.138, NK −0.125), whereas B cells, dendritic cells, megakaryocytes and FCGR3A+ monocytes had positive residuals, from +0.079 to +0.152; the probability of this split when three of seven residuals are negative at random is 1/35 = 0.029. The standard deviation of the fifteen residuals was 0.096, and the largest residual was that of stimulated megakaryocytes (+0.152).

The relation is driven mainly by the spread between conditions. Among the seven unstimulated conditions, whose measured overlaps lie between −0.12 and +0.34, the rank correlation between model value and measured overlap was 0.429 (p = 0.34). The calibration was not part of the analysis plan and is exploratory (Additional file 1, Table S1).

### Stimulus-specific eQTL

We next asked whether the same pair of conditions identifies genetic effects that depend on stimulation. Fine-mapped eQTL from two monocyte stimulation studies [5, 6] were taken from the eQTL Catalogue [18]. A variant–gene pair was labelled activation-specific or resting-specific according to the condition in whose credible set it appeared, and was kept only if its absolute z score in the other condition was below 2 (see “Condition-specific labels from credible-set membership” below). For each site we computed the log ratio of the absolute predicted effect under the stimulated condition to that under the resting condition, and compared activation-specific with resting-specific sites using a two-sided Mann–Whitney test among sites whose larger predicted effect exceeded 0.1.

The analysis plan named the Fairfax interferon-γ arm as the main analysis. It was inconclusive: 10 activation-specific against 23 resting-specific sites, p = 0.096, difference in group medians +0.37 (95% CI −0.20 to +1.54). Neither Fairfax LPS arm, which the plan listed for comparison, separated the groups (24 h: p = 0.44; 2 h: p = 0.34), and both point estimates had the opposite sign to that of the interferon-γ arm. An earlier run of the main analysis with credible-set labels, made before version 2 of the plan was written, gave p = 0.0489 by rank sum and p = 0.0383 by the permutation test specified in version 1, with no difference in a placebo split (p = 0.276). It would not have met the plan’s criterion for support, which also required a clearly weaker result in the LPS 2-h arm. In that run the LPS 2-h arm showed a difference larger than that of the interferon-γ arm and opposite in sign (median log2 ratio −0.808 at activation-specific against −0.301 at resting-specific sites); the interferon-γ difference had the sign expected in version 1, but it arose from smaller predicted effects under stimulation at resting-specific sites, not from the larger effects at activation-specific sites that version 1 had expected.

The plan treated the four Quach arms as replications and expected the largest difference for influenza; this part of the plan was written before any prediction for a Quach site existed (Methods). Influenza separated the groups (12 against 18 sites, p = 0.0019; 0.0076 after correction for the four replication arms and 0.0133 for all seven arms). LPS, R848 and Pam3CSK4 did not (p = 0.45, 0.74 and 0.95), with differences in group medians between −0.002 and +0.08. Arms within a study share sites: within the Quach study, LPS and Pam3CSK4 share 3 of the 6 activation-specific and 4 of the 7 resting-specific sites of the LPS arm, and the two Fairfax LPS arms share 16 of the 26 resting-specific sites of the 24 h arm (Additional file 1, Table S4b), so a correction by the number of arms is conservative. A criterion added to the plan later (supplement D) would have set aside an arm whose transcriptional response was negatively correlated with the interferon-β response; no arm met it (match scores +0.086 to +0.621; Methods), so the reversed Fairfax LPS arms count as directional inconsistencies. Counts, medians, confidence intervals, leave-one-out values and results under a gate on the smaller term are given for every arm in Additional file 1, Table S4.

### Source of the influenza signal

The influenza result was robust to leave-one-out (worst p = 0.00374), to placebo splits (p between 0.28 and 0.94), to removing the three sites shared with the Fairfax study (11 against 16 sites, p = 0.0072) and to a stricter label threshold (12 against 17 sites, p = 0.0015). Per-site medians of the bounded index were +0.478 for activation-specific and −0.134 for resting-specific sites.

The visibility gate in the plan, however, constrains only the larger of the two predicted effects. For most activation-specific sites the ratio exceeds 1, so the resting-condition effect is the smaller term and is not constrained. Seven of the twelve activation-specific sites had resting-condition predictions below 0.1 (median 0.077), compared with two of eighteen resting-specific sites (median 0.172). Requiring both effects to exceed a threshold removed the difference (Table 2), although part of this loss reflects the smaller sample. Testing the two terms separately does not depend on sample size: the groups did not differ in the stimulated-condition prediction (p = 0.144) but did differ in the resting-condition prediction (p = 0.00373; Fig. 3). The influenza result therefore reflects a difference in resting-condition baselines rather than stimulation dependence.

**Figure 3:**
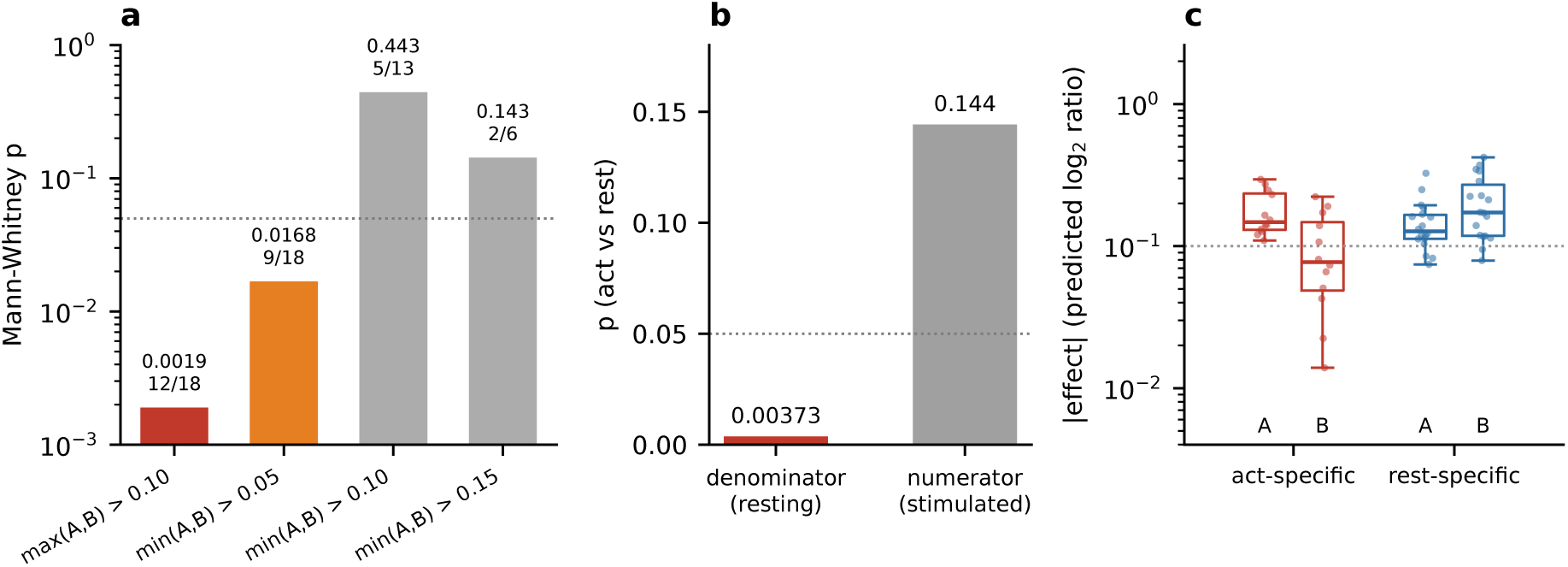
Decomposition of the site-level association in the influenza arm. (a) Mann–Whitney p for the activation-specific versus resting-specific contrast under four visibility gates. The gate on the larger of the two effects, as specified in the plan (max(A,B) > 0.1), gives p = 0.0019; requiring both effects to exceed 0.1 gives p = 0.443 (min(A,B) > 0.1, n = 5/13). (b) The same contrast split into its two terms: the groups differ in the resting-condition effect (p = 0.00373) but not in the stimulated-condition effect (p = 0.144). (c) Distributions of the stimulated (A) and resting (B) effect magnitudes by group; the activation-specific group has the smaller resting-condition effects.

**Table 2.** The site-level contrast under four visibility gates, in the influenza arm.

| Gate | n (activation / resting) | rank-sum p |
|---|---|---|
| max(A,B) > 0.10, as specified in the plan | 12 / 18 | 0.0019 |
| min(A,B) > 0.05 | 9 / 18 | 0.0168 |
| min(A,B) > 0.10 | 5 / 13 | 0.443 |
| min(A,B) > 0.15 | 2 / 6 | 0.143 |

### Control analyses

The first-generation model, which has no stimulated-state tracks, also separated the two groups by baseline effect size in its CD14+ monocyte RNA track (median absolute predicted effect 0.037 for activation-specific against 0.082 for resting-specific sites, p = 0.044), but not in its DNase track (p = 0.28). The two groups therefore differ in a property unrelated to conditioning. Conditioning on a mismatched cell type (CD4 T or B cells) collapsed the predictions for activation-specific sites to median absolute values of 0.006 and 0.021, with seven of twelve below 0.01, so the absence of separation in this control (p = 0.92 and 0.82) is uninformative. Because this control changes the cell type rather than the state, it shows only that the output depends on the conditioning input.

### Condition-specific labels from credible-set membership

The site-level analysis depends on how condition specificity is defined. A variant that appears in the credible set of one condition but not the other is often treated as condition-specific, but a difference in significance is not itself a significant difference [19], and joint analyses of eQTL across tissues find that most effects are shared, often differing in magnitude rather than presence [20]. We therefore looked up each labelled variant in the other condition’s full summary statistics. Of the rows labelled condition-specific by credible-set membership, 1,077 of 1,506 in Fairfax (71.5%) and 2,754 of 3,628 in Quach (75.9%) had an absolute z score of at least 2 in the other condition (Table 3; Fig. 4). A further 18 and 34 rows could not be tested because the variant was absent from the other condition, and 2 and 421 because the gene or probe had no record there; these rows were excluded and not treated as evidence of no effect. Lowering the threshold to 1.9 moved 15 Fairfax rows and 27 Quach rows and changed no conclusion. Under the looser labels each arm had two to three times as many sites, and the site-level estimates changed with every reasonable change of definition.

**Figure 4:**
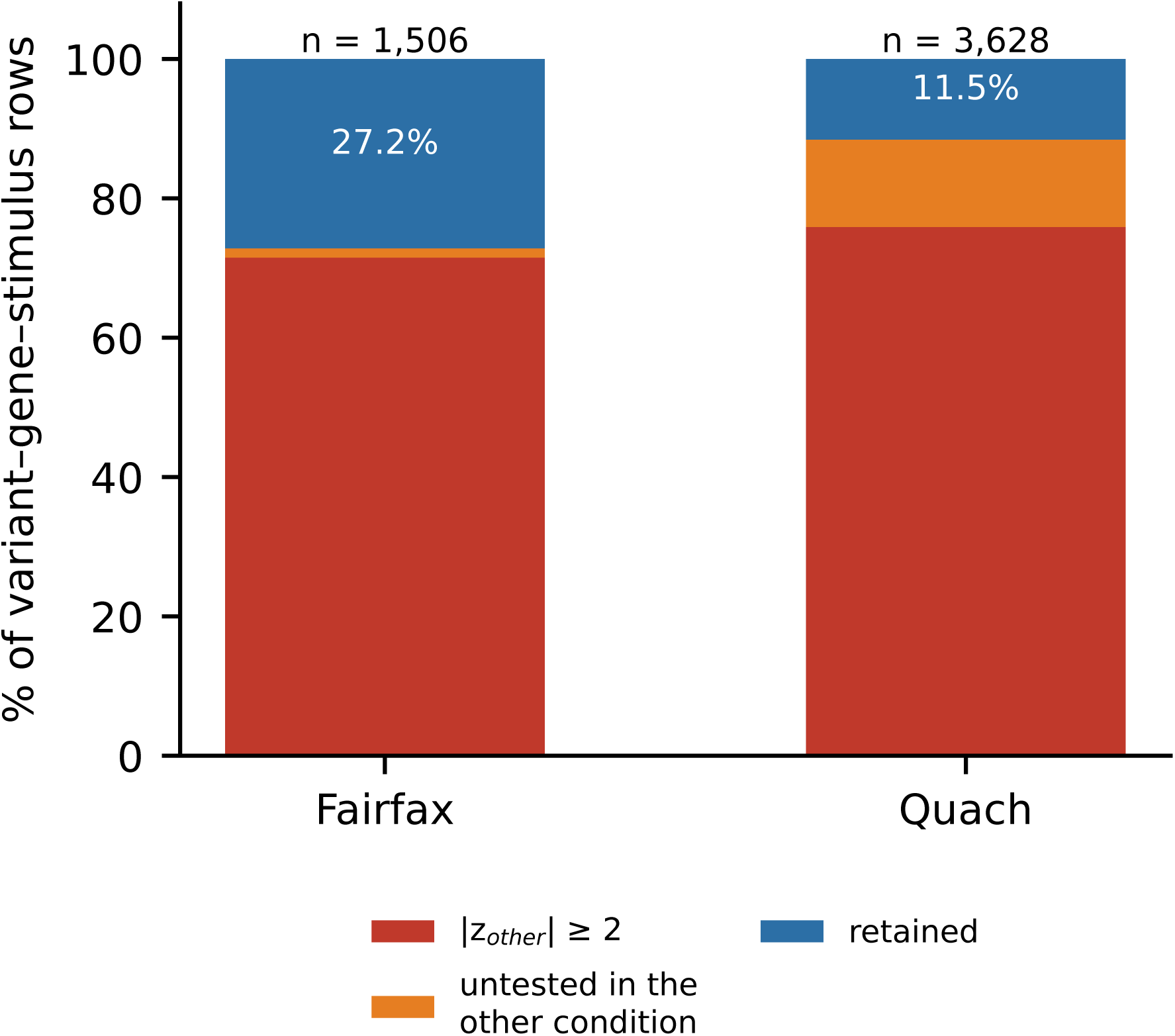
Attrition of credible-set condition-specific rows under the strict definition. Fraction of variant–gene–stimulus rows removed when the strict definition is applied: rows whose effect in the other condition reaches |z| = 2, rows untested in the other condition (variant or probe for Fairfax, variant or gene for Quach), and rows retained after both exclusions. Fairfax: 1,077 of 1,506 condition-specific rows removed (71.5%); Quach: 2,754 of 3,628 (75.9%).

**Table 3.** Attrition of credible-set condition-specific rows under the strict definition.

|  | Fairfax | Quach |
|---|---|---|
| Condition-specific label rows (activation + resting) | 1,506 | 3,628 |
| of which validation comparisons | 1,506 | 3,627 |
| removed: absolute z ≥ 2 in the other condition | 1,077 (71.5%) | 2,754 (75.9%) |
| removed: variant not tested in the other condition | 18 | 34 |
| removed: gene or probe not tested in the other condition | 2 | 421 |
| retained: activation / resting | 146 / 263 | 219 / 200 |
| Shared rows, excluded from the comparison | 420 | 1,751 |
| All label rows (1,506 + 420; 3,628 + 1,751) | 1,926 | 5,379 |
| Shared rows with opposite effect direction | 4 | — |

## Discussion

Conditioning AlphaGenome 2 on a single-cell profile of interferon-β-stimulated monocytes produced expression predictions that tracked the measured response, and the predicted response scaled with the similarity between each conditioning state and the target state. In contrast, we found no reliable evidence that the same conditioning identifies stimulus-specific eQTL: the main analysis was inconclusive, and the only significant arm reflected a difference in resting-condition baselines. The model therefore reproduced a state-level expression response, whereas its ability to capture state-dependent variant effects remains unestablished.

### Expression-level prediction

Our expression-level result is not directly comparable with benchmarks of perturbation-response models such as scGen [9], which predict the response of a held-out cell type from the responses observed in other cell types of the same dataset. Here the model received the stimulated state as input and predicted expression from sequence, although none of its built-in conditions is labelled with this stimulus. Whether a comparably simple baseline would perform as well, as reported for the prediction of genetic perturbation effects [12], remains to be tested. The calibration indicates that the predicted response was attenuated by about one quarter on average and that the attenuation was not uniform: stimulated T and NK states were consistently under-predicted, so users conditioning on such states should expect the response to be understated. Because the reliability of the measured axis was 0.983, the analysis is limited mainly by the number of conditions.

### Site-level result and its possible explanations

Neither catalogue contains an entry labelled with an experimental treatment. Across 4,430 first-generation tracks and 2,799 second-generation conditions, keyword searches over every text field returned no cytokine, Toll-like receptor ligand, virus, glucocorticoid or androgen treatment under either a narrow or a wide keyword list (Additional file 1, Table S3). The only matches were two GTEx skin labels (sun-exposed and not sun-exposed) and one GTEx label for Epstein-Barr-transformed lymphocytes, none of which is an experimental treatment under our definition (Methods). Lymphoblastoid lines appear under their cell-line names; GM12878 and the lines of its parents (GM12891 and GM12892) alone contribute 131 tracks in the first-generation catalogue. The catalogues do carry cell states. Under the wide list (activated, activation, effector, memory), 29 first-generation tracks and 135 second-generation conditions are labelled with a differentiation or activation state, such as memory and effector memory T cells, memory B cells, activated T cells and activated fibroblasts. Memory and effector labels occur in both catalogues, whereas the word activated occurs only in the second-generation conditions.

The absence of a labelled treatment is one possible explanation for the site-level result, not a demonstrated cause. The same result is expected if sequence-to-expression models predict variation between genes better than variant effects between individuals [13, 14]; if the model underestimates the effects of causal eQTL variants, as reported for the first-generation AlphaGenome [21]; or if the conditions do not match, since none of the eQTL arms used interferon-β. The site-level analysis does not show that the conditioning carries no information about variant effects. It shows that we could not detect any in two studies and seven stimulus arms once both conditions were required to have a measurable predicted effect, with at most a few dozen sites per arm. A ground truth based on allelic activity measured directly in both conditions, such as reporter assays run under two treatments [22–24], would test the same question with an order of magnitude more sites.

### Implications for evaluating condition-dependent predictions

Two issues in our site-level analysis are likely to affect other evaluations of condition-dependent predictions. First, labelling eQTL as condition-specific by credible-set membership retained mostly variants with significant effects in both conditions; condition specificity should instead be defined by testing the effect in both conditions or by modelling the conditions jointly [20]. Second, filtering a ratio of two predicted effects on the larger term leaves the denominator unconstrained, so groups that differ in how often their denominators lie near the detection floor separate without any difference in condition dependence. Problems with ratio variables are long recognised [25]; in our data this one produced p = 0.0019 in the arm that the plan expected to show the largest difference. Three practices would have detected it: filtering on the smaller of the two terms, reporting the two terms separately for each group, and testing each term against the grouping. Effect sizes are also better reported in bounded form. The log ratio for activation-specific sites in the influenza arm is +1.504, which reads as a 2.8-fold difference, whereas the bounded index, the difference of the two terms over their sum, is +0.478 and cannot be inflated by a small denominator; per site the two are monotone transforms of each other, so rank-based tests give identical p values. The same issue arises whenever predicted quantities are compared between two conditions, cell types or model versions by taking their ratio.

### Related work and future directions

A literature search on 24 November 2026 found three independent evaluations of the first-generation model [15, 16, 26], one fine-tuning of it [27], and no evaluation of cell-state conditioning. Our site-level result is consistent with the finding that annotations derived from these models do not improve common-variant discovery beyond standard functional annotations [16], although the two studies address different questions. An independently developed cell-state-conditioned model with its own variant-effect scoring [3] was not evaluated here; both analysis issues described above apply to it. Further work should test the expression-level result in other datasets, stimuli and cell types, use allelic activity measured in paired conditions as ground truth, build conditions from experimentally perturbed single-cell data, and compare conditioned models.

### Limitations

The expression-level evidence comes from one cell type, one stimulus, one dataset and 119 genes, all on chromosomes 1 to 7 and X; the fifteen calibration conditions come from the same dataset and are correlated against the same measured response, so they are not independent replications. The conditioning and measurement cells share eight donors, all patients with systemic lupus erythematosus, a disease often accompanied by a type I interferon signature in blood; if the unstimulated cells carry such a signature, the measured interferon-β fold changes would be compressed. The calibration slope is an average: individual conditions deviate from the fit by up to 0.15, and the calibration was added after the analysis plan. The site-level analysis is underpowered in every arm: strict labelling leaves at most a few dozen visible sites per arm, some with only 6 or 7 per group and, under the gate on the smaller term, as few as two sites per group (Additional file 1, Table S4). The mismatched-condition control is uninformative because the predictions collapse under it. Finally, the statement that neither catalogue contains an entry labelled with an experimental treatment is limited to the two keyword lists and the fields searched; because the catalogues label activation by cell state rather than by how the state arose, some activated states may come from experimentally stimulated cells. Further technical notes, including sensitivity to the pseudocount and the overlap between the two eQTL studies, are given in Additional file 1, Supplementary Note 2.

## Conclusions

Conditioning AlphaGenome 2 on a single-cell profile of a stimulated state reproduced the expression response of monocytes to interferon-β, and the predicted response scaled with the similarity of the conditioning state to the target. We found no reliable evidence that the same conditioning identifies stimulus-specific eQTL; the one significant result arose from an analysis artifact. Evaluations of conditioned models should assess state-level expression and state-dependent variant effects separately, define condition-specific eQTL by testing both conditions, require measurable effects in both conditions when comparing them, and report every planned comparison.

## Methods

### Single-cell data

All expression data come from GSE96583 batch 2 [17]: peripheral blood mononuclear cells from eight donors with systemic lupus erythematosus, assayed unstimulated and after six hours of interferon-β stimulation, with cell-type annotations and singlet calls from the series. Only singlets were used. Counts were pooled within each cell type and library and converted to counts per million (CPM); pooling, rather than treating cells as replicates, avoids the inflated false-discovery rates that follow from the latter [28].

### Conditioning and scoring

Each condition was built from 256 cells, the maximum accepted by the cell encoder, drawn once per library with default_rng(7) without replacement; all cells were used where a population had fewer than 256 (megakaryocytes, 122 and 134; dendritic cells, 203 and 228). Predictions were made with the denoised single-cell gene-expression head under a 200 kb mean mask.

The 40 input intervals were the 1,048,576-bp bins containing the most genes among the first 1,500 expressed genes in the order of the count matrix; each interval was centred on its bin midpoint. The count matrix orders chromosomes by length (1 to 7, then X, then the rest), so these genes, and therefore all 40 intervals, lie on chromosomes 1 to 7 and X. Gene midpoints, (start + end)/2, assigned genes to bins, whereas the window-clustered bootstrap grouped genes by the Ensembl annotation start, which for a gene on the minus strand is its 3 end. The deposited predictions cover the 246 genes in these intervals that have a measured value; 119 of them pass the activity filter, falling in 35 of the 40 intervals.

The analysis set was defined once, on the matched condition only, as genes whose larger predicted activity across the two states exceeds 1 and that have a measured value in the count matrix. With a pseudocount of 1 the measured log ratio is always finite, so in practice the activity threshold selects the set. This gives 119 genes in 35 windows, which are used for every condition; defining the set per condition would give 116 to 132 genes and would not be comparable across conditions. The predicted log fold change is log2 of (treated activity + 1) over (baseline activity + 1), and the measured response is log2 of (stimulated CPM + 1) over (control CPM + 1).

### Cell sets used for each quantity

The measured response and the measured overlap use different cell sets (Table 4). The measured response excludes the 256 conditioning cells in each library, so that it shares no cells with the conditioning input. The measured overlap describes cell state and uses all cells of each type. The control library contains 2,785 CD14+ singlets and the stimulated library 2,590, so the two measured-response sets differ in size after 256 cells are removed from each.

**Table 4.** Cells used for each quantity.

| Quantity | Cells | Library (counts) |
|---|---|---|
| Measured response, control | 2,529 CD14+ monocytes, the 256 conditioning cells excluded | 5,634,862 |
| Measured response, stimulated | 2,334 CD14+ monocytes, the 256 conditioning cells excluded | 5,762,086 |
| Measured overlap, baseline | 2,785 CD14+ monocytes, none excluded | 6,192,051 |
| Measured overlap, numerators | 122 to 5,161 cells per type, none excluded | see Additional file 1, Table S2 |

### Gene hold-out

For the hold-out, the 119 analysis genes were deleted as columns from the single-cell matrix supplied to the encoder, reducing it from 35,635 to 35,516 genes, and both conditions were rebuilt and rescored. No other input changed.

### Measured overlap and calibration

For each cell population, the measured overlap is the Spearman correlation between log2 of (that population’s CPM + 1) over (control CD14+ monocyte CPM + 1) and the measured interferon-β response. The model value is the same correlation with the predicted log fold change in place of the measured one. Unstimulated populations use the control library and stimulated populations the stimulated library; the matched condition’s measured overlap is 1 by construction.

The calibration is an ordinary least-squares fit of model value on measured overlap over the fifteen points. The reliability of the measured axis was estimated by splitting each cell type’s cells in half, computing baseline and numerator each from its own half, and correlating the two resulting vectors across the fourteen non-matched conditions; over 20 repetitions the half-sample reliability was 0.967 (SD 0.018), giving a Spearman–Brown full-sample reliability of 0.983. The disattenuated slope is the fitted slope divided by that value.

### Uncertainty and null distributions

Confidence intervals are window-clustered bootstraps: windows were resampled with replacement, rows concatenated and the statistic recomputed (2,000 draws, seed 0). A gene-level bootstrap would understate the uncertainty because genes within a window are correlated; both approaches understate it relative to resampling the conditioning cells. The split-half null halves the control cells, builds a condition from each half and scores the resulting log ratio against the measured response (50 draws). In one draw, predictions were available for only 62 of the 119 analysis genes, because scoring failed for part of one half’s intervals and only genes scored in both halves enter a draw; that draw is included in the reported distribution, and excluding it leaves none of the remaining 49 draws at or above the observed value. The depth comparison binomially thins the disjoint counts to the depth of the earlier shared-cell measurement (20 draws) and compares the resulting correlation with the shared-cell value at that depth.

### Site-level analysis

Fine-mapped eQTL were taken from eQTL Catalogue release 8 [18] for two monocyte studies [5, 6], fine-mapped with SuSiE [29]. A site is activation-specific if it enters only the stimulated condition’s credible set, resting-specific if it enters only the resting one, and shared if it enters both. Under the strict definition used throughout, a site additionally requires an absolute z score below 2 in the other condition’s full summary statistics; sites not tested there are counted separately (variant, gene or probe not tested) and are not treated as absence of effect, and shared sites require the same effect direction in both conditions.

The z scores for the other condition were retrieved from the remote tabix-indexed summary statistics. For the same variant, beta is identical in the credible-set file and in the nominal summary statistics of a study, but the standard error in the credible-set files is systematically smaller (median 1.55% in Fairfax and 3.24% in Quach), so the two files do not carry interchangeable z scores. The validation criterion written before the lookup, agreement within 0.01, therefore failed for 1,482 of 1,506 Fairfax and 3,617 of 3,627 Quach comparisons. After the labels had been built and the first group comparison made, we replaced it with sign agreement of 100%, a rank correlation above 0.999 and at least 99.5% agreement on the absolute-z-below-2 classification; these thresholds were set after the validation results were known. Under them, Fairfax agreed on 100% of 1,506 comparisons (rank correlation 0.99996) and Quach on 99.945% of 3,627, with the two disagreements at absolute z of 1.898 and 2.020. Twenty sites not found in the summary statistics were rechecked with a wider window, and no missed records were found. The revision changed how the lookup was validated, not which sites the labels contain.

The test statistic is log2 of the ratio of absolute predicted effects, stimulated condition over resting condition. Following the plan, arm-level comparisons use a visibility gate on the larger of the two absolute effects (> 0.1); because that gate leaves the smaller term unconstrained, results under a gate on the smaller term are reported for the influenza arm in Results and for all seven arms in Additional file 1, Table S4. Effect sizes are reported as the bounded index, the difference of the two terms over their sum. All medians of site-level quantities are per-site medians: for each site the log ratio and the bounded index are computed, and the median of those values is reported. Group comparisons used two-sided Mann–Whitney tests. Robustness was assessed by leave-one-out, placebo splits of the control condition, removal of sites shared between the two studies, and a stricter label threshold.

The match score, used only for the criterion added in supplement D, is the Spearman correlation across genes between an arm’s transcriptional response and the interferon-β response in the Kang data, log2((CPM stimulated + 1)/(CPM control + 1)) over all CD14+ monocytes. An arm’s response is the log2 ratio of median TPM between the stimulated and naive conditions for the Quach arms; for the Fairfax arms it is the median over donors of the difference between stimulated and naive microarray log intensities, taken per probe and then as the median over the probes of a gene. Genes enter if they carry an eQTL label for that arm, have median TPM above zero in both conditions (Quach) or are present in both sources (Fairfax), and have a row in the pooled CD14+ count matrix, which leaves 555 to 757 genes per arm; 95% intervals come from resampling genes with replacement (1,000 draws).

### Catalogue keyword searches

Counts of conditions and tracks carrying a given label were computed over the full returned tables, 2,799 conditions and 4,430 tracks, by concatenating every text field and matching case-insensitively. Two keyword lists, a narrow and a wide one, are reported; both lists and the fields searched are given in Additional file 1, Table S3. We counted as an experimental treatment a stimulus, drug or infection applied to cells before profiling; the derivation of a cell line, including EBV transformation or reprogramming to pluripotency, is not a treatment. The genetically modified tracks in the first-generation catalogue (734, all TF ChIP-seq) carry a tag on the factor under study, introduced by CRISPR insertion at the endogenous locus (694) or by stable transfection of a tagged construct (40), to identify where the factor binds rather than to perturb the cell; we did not count them as treatments.

### Analysis plan and deviations

The analysis plan was kept in versioned files with a local SHA-256 hash chain; there was no external registration or timestamp, so all timings rest on local file records. Version 1, written on 13 November 2026, named the Fairfax interferon-γ arm as the main analysis, to be tested as a mean difference of log ratios by permutation. Version 2 was written on 14 November, after the main analysis had been run once with credible-set labels (p = 0.0489 by two-sided rank sum). It replaced the main test with a two-sided rank sum, because a recomputation had shown that the mean difference was dominated by sites whose denominator approached zero; adopted the strict label definition and the 0.1 visibility threshold; and added the four Quach arms as replications, expecting influenza to show the largest difference, before any Quach prediction existed. It also rewrote the expected pattern to match what had been seen in the Fairfax arm: version 1 expected larger predicted effects under stimulation at activation-specific sites, whereas version 2 expected smaller ones at resting-specific sites; the expected sign of the group difference was the same. In this paper, “the plan” refers to version 2 together with its first supplement (supplement A). The plan required, for support of the main analysis, a significant main test (p < 0.05), a placebo split showing no difference, and a clearly weaker result in the LPS 2-h arm; it stated no rule for the replication arms.

Changes made after the plan are reported as such. Supplement C replaced supplement A’s rule for multiple microarray probes with exact matching on the molecular trait identifier, after the Quach predictions existed but before any strict label was built or any Quach comparison was run. Supplement D added the mismatch criterion described above; it was written after the Fairfax results under credible-set labels were known and before the strict labels were built or any Quach comparison was made, and no arm met it. The z-score validation criterion was replaced after the labels had been built and after the first group comparison (see above). The expression-level analyses were run before version 1 and the calibration was added later; both are exploratory. Additional file 1, Table S1 gives the status of each reported analysis relative to the plan, and Supplementary Note 1 gives the history of the plan with times; the complete log of changes is deposited with the code (`provenance_log.csv`).

### Use of AI-assisted tools

DeepSeek-V4.1-Flash was used as a coding and analysis agent under the direction of the authors, who decided which analyses to run and approved each reported result; analyses added after the analysis plan are identified in Additional file 1, Table S1. It wrote and ran the scoring and analysis code, plotted the figures from the archived tables and typeset the manuscript (13–16 November 2026), and corrected the text and reran analyses during revision (22–24 November 2026). A co-author recomputed the numbers reported in the version of 16 November independently of the analysis code, using only the archived per-gene tables, with Anthropic Claude Opus 5 and earlier model versions (13–16 November 2026). Before submission, an automated script (`audit_numbers.py`) checked 72 printed values against the deposited tables and run logs and found no discrepancies. Both models were used to draft and revise text (15–16 November 2026), and Anthropic Claude Opus 5.5 was used to review the manuscript and edit the text during revision (22–24 November 2026); the authors reviewed and edited all such text. No figure was generated as an image, no model is an author, and the authors take full responsibility for the content.

## List of abbreviations

API: application programming interface
ChIP-seq: chromatin immunoprecipitation followed by sequencing
CI: confidence interval
CPM: counts per million
EBV: Epstein–Barr virus
eQTL: expression quantitative trait locus or loci
GTEx: Genotype-Tissue Expression project
GWAS: genome-wide association study
IFN: interferon
LPS: lipopolysaccharide
NK: natural killer
PBMC: peripheral blood mononuclear cell
SDK: software development kit
SuSiE: sum of single effects
TF: transcription factor
TPM: transcripts per million.

## Declarations

### Ethics approval and consent to participate

Not applicable. All data analysed here are public and were previously collected under the original studies’ approvals; no human participants were recruited for this work.

### Consent for publication

Not applicable.

### Availability of data and materials

All primary data are public. Single-cell expression data are from GEO series GSE96583 [17], and fine-mapped eQTL from eQTL Catalogue release 8 [18]. Model predictions were obtained through the public API (alphagenome Python SDK v2.0.0) between 13 and 16 November 2026; the AlphaGenome 2 technical report accessed on 16 November has since been superseded by a preprint [4].

The derived tables are deposited so that every number in this paper can be recomputed without access to the model. The deposit contains the per-gene predictions for all fifteen conditions, cell-type CPM including the two columns that define the measured response, the cell counts and library sizes behind every column, the reliability vectors, the calibration summary, the site-level tables for all seven arms (labels, z scores and predicted effects under both conditions), the run log check_fixed.log, the full first-generation track table (v1_all_tracks.csv, 4,430 rows), the full second-generation condition table (`ag2_conditions.csv`, 2,799 rows) and the complete log of changes made during the independent recomputation and revision (`provenance_log.csv`). The README states two conventions that are easy to get wrong: the columns that define the measured response differ from the CD14+ monocyte columns used for the measured overlap (using the latter gives 0.619 rather than 0.615 for the main result), and the analysis gene set must be derived from the matched condition’s rows alone.

Code for the analysis and for the independent recomputation is available at https://github.com/hhdgit0149/alphagenome2-conditioning-evaluation. The code, the derived tables and the manuscript sources used for this paper are archived together at https://doi.org/10.5281/zenodo.24494903 (v1.1.0). The code is released under the MIT licence and the other files under CC BY 4.0, except that files containing AlphaGenome outputs, or derived from them, remain subject to the AlphaGenome Output Terms of Use. The two codebases were written separately from the same specification; discrepancies between them that changed a reported number are recorded in `provenance_log.csv`. An earlier version of this manuscript, under a different title, was posted as a preprint at bioRxiv (https://doi.org/10.64898/2026.11.16.767041) on 17 November 2026 under a CC BY 4.0 licence.

### Authors’ contributions

HH: conceptualization, methodology, software, formal analysis, investigation, data curation, visualization, writing – original draft. JS: conceptualization, methodology, software, formal analysis, investigation, data curation, visualization, writing – original draft. JX: validation, investigation, writing – review and editing. MA: validation, investigation, writing – review and editing. YY: validation, investigation, writing – review and editing. YH: validation, data curation, writing – review and editing. ZW: conceptualization, supervision, project administration, writing – review and editing. QX: conceptualization, supervision, project administration, writing – review and editing. SW: conceptualization, supervision, project administration, writing – review and editing. All authors read and approved the final manuscript.

## Acknowledgements

We thank the investigators of the Kang, Fairfax and Quach studies and the eQTL Catalogue for making their data publicly available. The use of AI-assisted tools is described in Methods.

## Supplementary information

**Additional file 1** (PDF). Supplementary Tables S1–S4b, Supplementary Figures S1–S2 and Supplementary Notes 1–2. Table S1, status of each reported analysis relative to the analysis plan; Table S2, cells and library sizes behind each expression column; Table S3, keyword lists, fields searched and counts; Table S4, site-level results for the seven stimulus arms; Table S4b, overlap of visible sites between arms within each study; Figure S1, effect of shared cells and sequencing depth on the expression-level correlation; Figure S2, split-half null and unmatched cell-type conditions; Supplementary Note 1, history of the analysis plan; Supplementary Note 2, additional technical notes.

