## Supplementary information (Tables S1-S3, Figures S1-S2) for "Cell-state conditioning in AlphaGenome 2 reproduces an unseen interferon response but shows no reliable stimulus-specific eQTL effects"

Contents: Table S1 (status of each reported analysis relative to the analysis plan); Table S2 (cells and libraries behind every expression column used in Figure 2); Table S3 (keyword lists, fields searched and counts for the catalogue analysis); Table S4 (seven stimulus arms); Table S4b (overlap of visible sites between arms); Figure S1 (effect of shared cells and sequencing depth); Figure S2 (split-half null and unmatched cell-type conditions); Supplementary Note 1 (history of the analysis plan); Supplementary Note 2 (additional technical notes).

### Table S1. Status of each reported analysis relative to the analysis plan

Table S1 lists each analysis reported in the main text and its relation to the analysis plan (Methods; Supplementary Note 1). Analyses run before plan version 1 precede the plan; analyses after plan version 2 were added later. Exploratory analyses were not used to test a hypothesis stated in the plan. The complete log of changes made during the independent recomputation and during revision, including corrections of reported values, is deposited with the code as `provenance_log.csv`. The relation to the plan refers to when an analysis was first run; later changes, including recomputation of reported values, are given in the Note column.

Status of each analysis reported in the main text. "Specified in plan version 1" and "Specified in plan version 2" refer to the hashed plan texts; "After plan version 2" includes later supplements, which are not part of the plan as defined in the main text. Dates are in November 2026; times are local (UTC+8).

| Analysis | Reported in | Relation to the plan | Exploratory | Note |
|---|---|---|---|---|
| Expression-level comparison (predicted vs measured log2 fold change, 119 genes) | Results; Table 1; Fig. 1a | Run before plan version 1 | Yes | Measured response recomputed on cells disjoint from the conditioning input on 15 November |
| Gene hold-out (all 119 analysis genes removed from the conditioning input) | Results; Table 1; Fig. 1b, c | Run before plan version 1 | Yes | All-gene version run on 15 November |
| Shared-cell versus depth-matched comparison | Results; Table 1; Fig. S1 | After plan version 2 | Yes | Run with the disjoint recomputation on 15 November |
| Split-half null | Results; Table 1; Fig. S2a | Run before plan version 1 | Yes | Extended from 8 to 50 draws and recomputed on the disjoint design on 15 November |

| Analysis | Reported in | Relation to the plan | Exploratory | Note |
| --- | --- | --- | --- | --- |
| Removal of the ten genes with the largest measured fold change | Results | After plan version 2 | Yes | Added during revision (23 November) |
| Calibration across fifteen conditioning states | Results; Fig. 2 | After plan version 2 | Yes | Twelve points first (15 November), extended to fifteen the same day |
| Main analysis: Fairfax interferon-$\gamma$ arm, strict labels, rank-sum test, gate on the larger term | Results; Table S4 | Specified in plan version 2 | No | Test, label definition and gate adopted after the main analysis had been run once with credible-set labels (p = 0.0489) |
| Main test as specified in version 1 (credible-set labels; permutation) | Results; Methods | Specified in plan version 1 | No | Replaced in version 2; p = 0.0383 |
| Fairfax LPS 2 h and 24 h comparison arms | Results; Table S4 | Specified in plan versions 1 and 2 | No | The plan required a clearly weaker result in the LPS 2-h arm |
| Four Quach replication arms; influenza expected to differ most | Results; Table S4 | Specified in plan version 2 | No | Written before any prediction for a Quach site existed |
| Strict label definition (absolute z below 2 in the other condition) | Results; Table 3; Fig. 4 | Specified in plan version 2 and supplement A | No | Rule for multiple probes replaced by exact matching on the molecular trait (supplement C, 14 November 10:50), before any strict label was built |
| Attrition of credible-set condition-specific labels (71.5%, 75.9%) | Results; Table 3; Fig. 4 | After plan version 2 | Yes | Follows from the strict label definition; denominator corrected on 22 November |
| Gate on the smaller of the two terms | Results; Table 2; Fig. 3a; Table S4 | After plan version 2 | Yes | Added on 15 November during the independent recomputation |
| Decomposition into stimulated-condition and resting-condition terms | Results; Fig. 3b | After plan version 2 | Yes | Added on 15 November during the independent recomputation |
| Mismatch criterion (match score) | Results; Methods; Table S4 | After plan version 2 (supplement D) | – | Written on 14 November at 20:15, after the Fairfax results under credible-set labels were known; no arm met it |

| Analysis | Reported in | Relation to the plan | Exploratory | Note |
|---|---|---|---|---|
| Validation criterion for the z-score lookup | Methods | After plan version 2 (supplement E) | – | Replaced on 14 November at 22:40, after the labels were built and the first Quach comparison was made |
| Controls: first-generation model and mismatched cell type | Results | After plan version 2 | Yes | Neither control appears in the plan text or in supplements A–E; both were added on 15 November during the independent recomputation |
| Catalogue keyword searches | Discussion; Table S3 | After plan version 2 | Yes | Full tables exported on 15 November; counts revised on 22 November |

### Table S2. Cells and libraries behind every expression column

The two axes of Figure 2 rest on different cell sets. The x axis (measured overlap) uses all singlet cells of each annotated cell type, because more cells only stabilise the estimate of that state. The measured interferon-β response, which enters the y axis, excludes in each library the 256 cells that built the conditioning input, so that the conditioning vector and the measured response share no cells. Cells are disjoint between the two sets; the eight donors are not (see Limitations in the main text).

Cells and libraries behind every expression column used in Figure 2. UMI per cell is library counts divided by cells used.

| Column | Role | Cells used | Library (counts) | UMI per cell | Cells excluded | Note |
|---|---|---|---|---|---|---|
| ctrl\|CD14+ Monocytes | x axis (CPM) | 2,785 | 6,192,051 | 2,223.4 | 0 | all singlet cells of this type; conditioning cells not excluded |
| ctrl\|CD4 T cells | x axis (CPM) | 5,161 | 6,131,014 | 1,188.0 | 0 | all singlet cells of this type; conditioning cells not excluded |
| ctrl\|CD8 T cells | x axis (CPM) | 1,082 | 1,069,128 | 988.1 | 0 | all singlet cells of this type; conditioning cells not excluded |
| ctrl\|NK cells | x axis (CPM) | 916 | 828,937 | 905.0 | 0 | all singlet cells of this type; conditioning cells not excluded |
| ctrl\|B cells | x axis (CPM) | 1,307 | 1,630,847 | 1,247.8 | 0 | all singlet cells of this type; conditioning cells not excluded |
| ctrl\|Dendritic cells | x axis (CPM) | 203 | 690,676 | 3,402.3 | 0 | all singlet cells of this type; conditioning cells not excluded |
| ctrl\|Megakaryocytes | x axis (CPM) | 122 | 205,612 | 1,685.3 | 0 | all singlet cells of this type; conditioning cells not excluded |
| ctrl\|FCGR3A+ Monocytes | x axis (CPM) | 739 | 1,838,222 | 2,487.4 | 0 | all singlet cells of this type; conditioning cells not excluded |

| Column | Role | Cells used | Library (counts) | UMI per cell | Cells excluded | Note |
|---|---|---|---|---|---|---|
| obs\|ctrl (CD14, disjoint) | y axis (observed response) | 2,529 | 5,634,862 | 2,228.1 | 256 | excluding the 256 conditioning cells from this library |
| stim\|CD14+ Monocytes | x axis (CPM) | 2,590 | 6,374,141 | 2,461.1 | 0 | all singlet cells of this type; conditioning cells not excluded |
| stim\|CD4 T cells | x axis (CPM) | 5,195 | 6,613,779 | 1,273.1 | 0 | all singlet cells of this type; conditioning cells not excluded |
| stim\|CD8 T cells | x axis (CPM) | 957 | 1,041,106 | 1,087.9 | 0 | all singlet cells of this type; conditioning cells not excluded |
| stim\|NK cells | x axis (CPM) | 1,067 | 1,071,317 | 1,004.0 | 0 | all singlet cells of this type; conditioning cells not excluded |
| stim\|B cells | x axis (CPM) | 1,256 | 1,649,925 | 1,313.6 | 0 | all singlet cells of this type; conditioning cells not excluded |
| stim\|Dendritic cells | x axis (CPM) | 228 | 723,162 | 3,171.8 | 0 | all singlet cells of this type; conditioning cells not excluded |
| stim\|Megakaryocytes | x axis (CPM) | 134 | 213,542 | 1,593.6 | 0 | all singlet cells of this type; conditioning cells not excluded |
| stim\|FCGR3A+ Monocytes | x axis (CPM) | 852 | 2,228,640 | 2,615.8 | 0 | all singlet cells of this type; conditioning cells not excluded |
| obs\|stim (CD14, disjoint) | y axis (observed response) | 2,334 | 5,762,086 | 2,468.8 | 256 | excluding the 256 conditioning cells from this library |

### Table S3. Keyword lists, fields searched and counts

Both keyword lists were applied to the full returned tables, not to a subset: 4,430 reference tracks for the first generation (AlphaGenome 1) and 2,799 built-in conditions for the second (AlphaGenome 2). The narrow list is ours; the broad list is an independent re-implementation. Every additional hit returned by the broad list was read by hand, and none describes an experimentally applied stimulus; the last column records that reading.

Keyword counts for the catalogue analysis (Discussion). Both lists were applied to the full returned tables: 4,430 reference tracks for AlphaGenome 1 and 2,799 built-in conditions for AlphaGenome 2. The last column records the manual reading of every additional hit returned by the broad list; none describes an experimentally applied stimulus.

| Dataset | Items | Keyword list | Keyword group | Hits | Hits adjudicated as a real perturbation |
|---|---|---|---|---|---|
| AlphaGenome 1 | 4,430 | broad | stimulus / cytokine | 3 | 0 |
| AlphaGenome 1 | 4,430 | broad | glucocorticoid | 0 | 0 |
| AlphaGenome 1 | 4,430 | broad | hormone | 0 | 0 |
| AlphaGenome 1 | 4,430 | broad | activated state label | 29 | 0 |
| AlphaGenome 1 | 4,430 | broad | prostate | 57 | 0 |
| AlphaGenome 1 | 4,430 | narrow | stimulus / cytokine | 0 | 0 |
| AlphaGenome 1 | 4,430 | narrow | glucocorticoid | 0 | 0 |
| AlphaGenome 1 | 4,430 | narrow | hormone | 0 | 0 |
| AlphaGenome 1 | 4,430 | narrow | activated state label | 0 | 0 |
| AlphaGenome 1 | 4,430 | narrow | prostate | 43 | 0 |
| AlphaGenome 2 | 2,799 | broad | stimulus / cytokine | 4 | 0 |
| AlphaGenome 2 | 2,799 | broad | glucocorticoid | 0 | 0 |
| AlphaGenome 2 | 2,799 | broad | hormone | 2 | 0 |
| AlphaGenome 2 | 2,799 | broad | activated state label | 135 | 0 |
| AlphaGenome 2 | 2,799 | broad | prostate | 23 | 0 |
| AlphaGenome 2 | 2,799 | narrow | stimulus / cytokine | 0 | 0 |
| AlphaGenome 2 | 2,799 | narrow | glucocorticoid | 0 | 0 |
| AlphaGenome 2 | 2,799 | narrow | hormone | 0 | 0 |
| AlphaGenome 2 | 2,799 | narrow | activated state label | 30 | 0 |
| AlphaGenome 2 | 2,799 | narrow | prostate | 19 | 0 |

#### Table S4. Seven stimulus arms

Site-level results for every arm. The larger gate requires the larger of the two absolute effects to exceed 0.1 and is the gate named in the plan; the smaller gate requires the smaller one. Confidence intervals are window-clustered bootstrap. The match score is the Spearman correlation between the arm's own transcriptional response and the interferon-β response in the Kang data (Methods). No arm met the mismatch criterion of supplement D.

| Study | Arm | n act / rest | Median difference (95% CI) | p, larger gate | Worst leave-one-out | p, smaller gate | n act / rest, smaller gate | Match score (95% CI) |
|---|---|---|---|---|---|---|---|---|
| Fairfax | IFN-γ 24 h | 10 / 23 | +0.367 (−0.196 to +1.536) | 0.096 | 0.19 | 0.12 | 6 / 17 | +0.389 (+0.313 to +0.461) |
| Fairfax | LPS 24 h | 9 / 26 | −0.187 (−0.880 to +0.762) | 0.44 | 0.74 | 0.7 | 7 / 18 | +0.086 (−0.002 to +0.171) |
| Fairfax | LPS 2 h | 7 / 32 | −0.583 (−1.330 to +0.770) | 0.34 | 0.63 | 0.33 | 2 / 23 | +0.258 (+0.169 to +0.341) |
| Quach | Influenza 6 h | 12 / 18 | +1.892 (+0.247 to +2.616) | 0.0019 | 0.00374 | 0.44 | 5 / 13 | +0.621 (+0.568 to +0.663) |
| Quach | R848 6 h | 13 / 16 | +0.018 (−0.713 to +0.767) | 0.74 | 1.0 | 1.0 | 5 / 10 | +0.426 (+0.364 to +0.486) |
| Quach | LPS 6 h | 6 / 7 | +0.077 (−0.962 to +1.824) | 0.45 | 0.76 | 0.79 | 3 / 5 | +0.286 (+0.214 to +0.353) |
| Quach | Pam3CSK4 6 h | 6 / 7 | −0.002 (−1.231 to +1.276) | 0.95 | 1.0 | 1.0 | 2 / 3 | +0.190 (+0.108 to +0.268) |

#### Table S4b. Overlap of visible sites between arms within each study

Number of visible sites shared by two arms of the same study, over the number in the first arm. Arms within a study are therefore not independent, and correcting by the number of arms is conservative.

| Study | Arm A | Arm B | Shared / arm A (activation) | Shared / arm A (resting) |
|---|---|---|---|---|
| Quach | Influenza 6 h | LPS 6 h | 1 / 12 | 3 / 18 |
| Quach | Influenza 6 h | R848 6 h | 2 / 12 | 5 / 18 |
| Quach | Influenza 6 h | Pam3CSK4 6 h | 1 / 12 | 2 / 18 |
| Quach | LPS 6 h | R848 6 h | 2 / 6 | 4 / 7 |

| Study | Arm A | Arm B | Shared / arm A (activation) | Shared / arm A (resting) |
|---|---|---|---|---|
| Quach | LPS 6 h | Pam3CSK4 6 h | 3 / 6 | 4 / 7 |
| Quach | R848 6 h | Pam3CSK4 6 h | 2 / 13 | 4 / 16 |
| Fairfax | IFN-$\gamma$ 24 h | LPS 24 h | 2 / 10 | 12 / 23 |
| Fairfax | IFN-$\gamma$ 24 h | LPS 2 h | 2 / 10 | 16 / 23 |
| Fairfax | LPS 24 h | LPS 2 h | 2 / 9 | 16 / 26 |

**Figure S1. Effect of shared cells and sequencing depth on the expression-level correlation**

Correlation between prediction and measurement for (i) the original analysis, in which the measured response was computed from the same 256 cells that built the conditioning input and the library was shallow; (ii) cells disjoint from the conditioning input after binomial thinning back to the original depth (mean of 20 thinnings; error bar, SD); and (iii) disjoint cells at full depth. The original value lies only slightly above the thinned disjoint value, so sharing cells did not measurably inflate the correlation; the lower value of the original analysis reflects its roughly ten-fold shallower library.

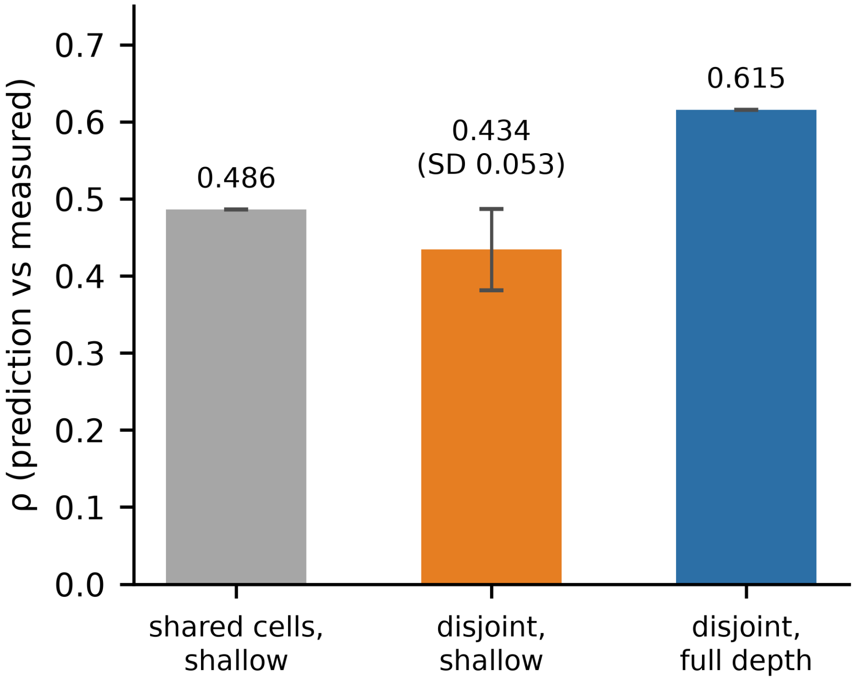


Figure S1. Effect of shared cells and sequencing depth on the expression-level correlation. Correlations between predicted and measured log2 fold change for the original analysis (same cells as the conditioning input, shallow library), for cells disjoint from the conditioning input after binomial thinning to the original depth (mean of 20 thinnings; error bar, SD), and for disjoint cells at full depth.

### Figure S2. Split-half null and unmatched cell-type conditions

(a) Split-half null: the control cells were randomly divided into two halves, each half was passed through the same pipeline as a condition, and the resulting correlation was compared with the measured response used throughout. None of 50 draws reaches the observed value (0 of 50; one-sided empirical p = 1/51 = 0.0196). One draw covers only 62 of the 119 genes (Methods). The null predictions are about an order of magnitude smaller than those of the matched condition. (b) The seven unstimulated cell-type conditions of Figure 2 (CD4 T, CD8 T, NK and B cells, megakaryocytes, dendritic cells and FCGR3A+ monocytes), plotted by the magnitude of their predicted change. Several produce predictions as large as or larger than the matched condition, yet their correlation with the measured response, which is their model value in Figure 2, lies between −0.12 and +0.30.

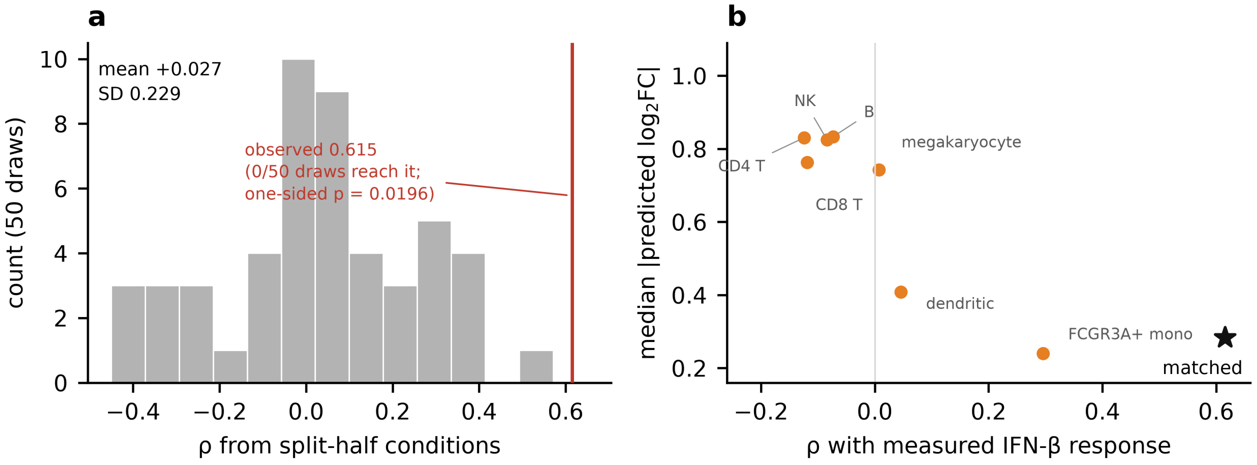


Figure S2. Split-half null and unmatched cell-type conditions. (a) Correlation from 50 split-half conditions, with the observed value marked. (b) Correlation with the measured response for seven unmatched cell-type conditions, plotted against the magnitude of their predicted change; the matched condition is the star.

### Supplementary Note 1. History of the analysis plan

The analysis plan was kept in versioned files with a local SHA-256 hash chain; there was no external registration or timestamp, so the timings below rest on local file records (all times UTC+8).

Version 1 was first hashed at 23:02 on 13 November 2026 and extended before it was hashed again at 23:19, with a further Fairfax scoring run at 23:11 in between; the 23:02 text is the first 105 lines of the file we hold, so what was added can be read off directly. It named the Fairfax interferon-$\gamma$ arm as the main analysis, tested as a mean difference of log ratios by permutation.

Version 2 was written at 00:50 on 14 November. It records the result of the main analysis under credible-set labels ($p = 0.0489$ by two-sided rank sum), computed one minute earlier; the rank-sum test, the strict label definition and the 0.1 visibility threshold were therefore adopted after that result was known. Version 2 replaced version 1's main test with a two-sided rank sum after a recomputation showed that the mean difference is dominated by sites whose denominator approaches zero. It also adopted the strict label definition as the main label definition, added the four Quach arms as replications with influenza expected to show the largest difference, and rewrote the expected pattern to match what had been seen in the Fairfax arm: where version 1 expected larger predicted effects under stimulation at activation-specific sites, version 2 expected smaller ones at resting-specific sites, with the same expected sign of the group difference. The Quach predictions did not yet exist.

The file of version 2 that we hold is the 00:50 text followed by five appended supplements, the last at 22:40 on 14 November. Hashing the file truncated at the start of each supplement reproduces the corresponding entry of the local hash chain, so the sections written at 00:50 were not edited afterwards; this check establishes content, not time, because the hash file has no external anchor. In the main text, "the plan" refers to the text hashed at 00:50 together with supplement A (01:20), both written before any prediction for a Quach site existed; analyses introduced by a later supplement are described as added after it.

The label rules were revised at 10:50 (supplement C), after the Quach predictions existed but before any Quach group comparison was run; supplement A's rule for multiple microarray probes was replaced at that point, before any strict label was built, by exact matching on the molecular trait identifier, and rows untested in the other condition are counted separately rather than treated as $|z| < 2$. The validation protocol written at 10:50 required agreement of the retrieved z scores with the credible-set values within 0.01 and stopping on any mismatch. Two validation runs, at 21:47 and 22:26, failed that criterion on almost every row because the standard errors in the two sources differ (Methods). We did not stop. The labels were built at 22:27 and the first Quach group comparison was made at 22:38. At 22:40 (supplement E) the validation criterion was replaced with sign agreement of 100%, a rank correlation above 0.999 and agreement of at least 99.5% on the absolute-z-below-2 classification; these thresholds were set after the validation results were known, and the revision changed how the lookup was validated, not which sites the labels contain.

Supplement D, appended at 20:15 on 14 November, after the Fairfax results under credible-set labels were known (00:49) and before the strict labels (22:27) or any Quach comparison (22:38), added the mismatch criterion: an arm whose match score is below zero, with a bootstrap interval entirely below zero, is marked uninformative and dropped from the cross-stimulus consistency check, and the verdict may not be used to turn a negative result into positive evidence or to explain why an arm performed well. No arm met it: the seven match scores run from +0.086 (Fairfax LPS 24 h, interval −0.002 to +0.171) to +0.621 (influenza), and the match score is +0.258 for the Fairfax LPS 2 h arm.

The expression-level analyses were run before either version and are exploratory; the calibration was added after the independent recomputation and is also exploratory. Table S1 gives the status of every result.

### Supplementary Note 2. Additional technical notes

**Interval width does not track cell number.** Megakaryocytes (122 control, 134 stimulated) and dendritic cells (203, 228) are the smallest populations, and their intervals are wide (unstimulated megakaryocytes 0.46, stimulated 0.43; unstimulated dendritic cells 0.39). But the three widest intervals belong to unstimulated NK, B and CD8 T cells, which rest on 916 to 1,307 cells (0.51, 0.49 and 0.49), and these three points, like the unstimulated megakaryocyte point, have measured overlaps close to zero, where the statistic is least stable. Cell number alone is therefore a poor guide to how well a point is determined.

**The split-half null bounds the p value.** With 50 draws the smallest attainable one-sided p is 1/51 = 0.020, and that is the value obtained; the null therefore bounds the p value rather than estimating it. Its predicted changes are also about an order of magnitude smaller than the real ones.

**Overlap between the two eQTL studies.** The two studies share 66 variant–gene pairs, 7.0% of the 944 unique pairs in the Quach study, which corresponds to 199 of its 2,170 retained rows; 3 of the 30 sites that drive the influenza arm are among them.

**Pseudocount convention.** Both sides use a pseudocount of 1. Varying it on the prediction side only, with the measured response held at 1, gives 0.606, 0.611, 0.616 and 0.614 at pseudocounts 0, 0.5, 2 and 5. Applying the same pseudocount to both sides gives 0.588, 0.609, 0.615, 0.621 and 0.647 at 0, 0.5, 1, 2 and 5; the zero case drops the five genes with a zero in the count matrix and rests on 114. The main comparison therefore lies between 0.588 and 0.647 across conventions, and no conclusion depends on the choice.

**Cell sets for the two axes of the calibration.** The measured overlap uses all singlets of each cell type, including the 256 used for conditioning, whereas the measured response excludes them. For the fourteen non-matched calibration conditions the conditioning cells are therefore a subset of those used for the measured overlap, and for populations with fewer than 256 cells they are the same cells; the effect of this sharing on the calibration was not tested. The measured axis and the measured response also share most of their CD14+ monocyte baseline, which deflates rather than inflates the fitted slope; recomputing against the disjoint baseline moves it from 0.728 to 0.731.

**The two attrition percentages are specific to our definition.** Our strict definition requires that the variant was tested in the other condition and had $|z| < 2$ there; a study that defines condition specificity by credible-set membership alone reports a different and larger set, and neither number can be substituted for the other.
